# Pharmacokinetics and Central Accumulation of Delta-9-Tetrahydrocannabinol (THC) and its Bioactive Metabolites Are Influenced by Route of Administration and Sex

**DOI:** 10.1101/2021.07.02.450963

**Authors:** Samantha L. Baglot, Catherine Hume, Gavin N. Petrie, Robert J. Aukema, Savannah H.M. Lightfoot, Laine M. Grace, Ruokun Zhou, Linda Parker, Jong M. Rho, Stephanie L. Borgland, Ryan J. McLaughlin, Laurent Brechenmacher, Matthew N. Hill

## Abstract

Up to a third of North Americans over 16 years old report using cannabis in the prior month, most commonly through inhalation. Animal models that reflect human cannabis consumption are critical to study its impacts on brain and behaviour. Nevertheless, most animal studies to date examine effects of cannabis through injection of delta-9-tetrahydrocannabinol (THC; primary psychoactive component of cannabis). THC injections produce markedly different physiological and behavioural effects than inhalation, likely due to distinctive pharmacokinetics of each administration route. The current study directly examined if administration route (injection versus inhalation), with dosing being matched on peak THC blood levels, alters the metabolism of THC, and the central accumulation of THC and its metabolites over time. Adult male and female Sprague-Dawley rats received either a single intraperitoneal injection of THC (2.5 mg/kg) or a single (15 min) session of inhaled exposure to THC distillate (100 mg/mL) vapour. Blood and brains were collected at 15, 30, 60, 90 and 240 minutes post-exposure for analysis of THC and metabolites through mass spectrometry-liquid chromatography. Inhalation results in immediate hypothermia, whereas injection results in delayed hypothermia. Despite achieving comparable peak concentrations of blood THC in both groups, our results indicate higher initial brain THC concentration following inhalation, whereas injection resulted in dramatically higher 11-OH-THC concentrations, a potent THC metabolite, in blood and brain that increased over time. Our results provide evidence that THC and its metabolites exhibit different pharmacokinetic profiles following inhalation versus injection, which could have significant impacts for data interpretation and generalizability. Accordingly, we suggest that translational work in the realm of THC and cannabis strongly consider using inhalation models over those that employ injection.

**Highlights:**

- Body temperature as well as blood and brain levels of THC and metabolites differ based on administration route
- THC inhalation results in immediate hypothermia, whereas THC injection results in delayed hypothermia
- THC inhalation results in higher initial brain THC levels than injection
- THC injection results in higher blood & brain 11-OH-THC levels than inhalation
- Translational cannabis work should strongly consider using inhalation over injection

## 1. Introduction

With recreational use of cannabis recently becoming legal in Canada and in some states across the U.S., as well as medicinal use being legal in many other countries, there is a growing need to better understand the effects of cannabis on the brain and behaviour. Up to a third of North Americans over 16 years of age report using cannabis in the prior month (Goodman et al., 2020), most commonly through pulmonary (i.e. inhalation) administration (Cuttler et al., 2016). Animal models provide an extremely valuable approach to studying the effects of cannabis, enabling control over composition, dose, and timing of exposure. Nevertheless, the majority of animal studies examine the effects of cannabis through parenteral (i.e. intraperitoneal [IP]) injections of delta-9-tetrahydrocannabinol (THC; the primary psychoactive component of cannabis) or cannabinoid receptor 1 (CB1R) agonists, which does not reflect a common route of human cannabis consumption nor reproduce the same physiological or behavioural effects of inhalation administration.

Inhalation of THC produces a much more rapid onset and offset of hypothermia, increases feeding behaviour, decreases locomotion, and fails to induce cross-sensitization to morphine in rats when compared directly to injected THC (Manwell et al., 2014; Nguyen et al., 2016). Further, while THC injections result in conditioned avoidance, inhalation produces a place preference (Manwell et al., 2014) and has reinforcing properties (as demonstrated by robust self-administration) in rats (Freels et al., 2020). The effects of THC inhalation on temperature, appetite, and locomotion in rodents are also typically shown in humans after exposure to cannabis smoke (Ashton, 2001; Kirkham and Williams, 2005; Sharma et al., 2012). The physiological and behavioural differences between inhalation and injections are likely due to the distinctive pharmacokinetic (PK) differences of each route of administration.

Following IP injection, compounds are absorbed primarily through the portal circulation and pass through the liver to undergo metabolism before reaching other organs, such as the brain (Lukas et al., 1971; Turner et al., 2011). Alternatively, inhalation provides rapid delivery of compounds into the blood stream, bypassing initial metabolism by the liver, and resulting in more immediate uptake by highly perfused tissues, such as the brain (Huestis, 2007, 2005; Turner et al., 2011). The amount and duration of compound absorption also differs, with injections delivering a single bolus versus inhalation delivering an ongoing infusion. As such, the PK profile of THC, specifically plasma THC concentrations, between injections and inhalation differ in timing, magnitude, and duration.

The most common form of human cannabis administration – inhalation (smoking or vaporization) – produces peak plasma THC concentrations 10-15 min after initial administration (Hartman et al., 2015; Huestis et al., 1992; Huestis and Cone, 2004; Newmeyer et al., 2016; Schwope et al., 2011; Spindle et al., 2019, 2018) with relatively rapid clearance of THC from plasma. In fact, plasma THC concentrations are only 15-20% of peak at 30 min following cannabis use, 8-10% at 60 min, and 2-3% at 180 min (Abrams et al., 2007; Newmeyer et al., 2016). However, as a result of individual differences in the number, duration, and spacing of puffs, as well as inhalation volume and hold time, the exact concentration of peak plasma THC in human studies is extremely variable. Peak plasma concentrations of THC range from 60-200 ng/mL following inhalation of cannabis flower (Abrams et al., 2007; Bidwell et al., 2020; Hartman et al., 2015; Huestis et al., 1992; Huestis and Cone, 2004; Newmeyer et al., 2016; Schwope et al., 2011), making animal models that can control dose and timing extremely valuable.

Rodent studies utilizing THC injections allow for control over both dose and timing; however, peak plasma THC concentrations are found at a slightly later timepoint following administration than with respect to inhalation (Javadi-Paydar et al., 2017; Nguyen et al., 2016; Taffe et al., 2020; Torrens et al., 2020). Further, clearance of THC from plasma following IP injections is much slower, with concentrations still roughly 65% of peak at 60 min and 50% at 120 min (Torrens et al., 2020). Rodent studies employing IP injections also utilize a wide range of dosages (3 to 20 mg/kg) producing an extensive span of peak plasma THC concentrations from 40-200 ng/mL (Javadi-Paydar et al., 2017; Nguyen et al., 2016; Taffe et al., 2020; Torrens et al., 2020), suggesting that they are comparable to the range seen in humans following cannabis use. Interestingly, brain THC concentrations following IP administration increase over time, peaking at 60-120 min following initial administration (Torrens et al., 2020).

Animal models utilizing vapour delivery of THC or whole cannabis extract have recently been validated (Javadi-Paydar et al., 2017; Manwell et al., 2014; Nguyen et al., 2016) and are able to control for dose and timing of exposure while also employing the most common route of cannabis consumption in humans. Several rodent studies have found plasma THC concentrations of 100-200 ng/mL following 30 min of exposure to 100-200 mg/mL of THC vapour (Javadi-Paydar et al., 2017; Nguyen et al., 2020, 2016; Taffe et al., 2020). Similar to human inhalation, plasma THC concentrations peak at around 15 min (Hložek et al., 2017) with relatively rapid plasma clearance as suggested by concentrations of ∼30% of peak at 60 min and 8-10% at 120 min (Javadi-Paydar et al., 2017; Nguyen et al., 2016; Taffe et al., 2020). Finally, opposite to injection, brain THC concentrations following inhalation peak at 15 min following initial administration and decrease over time (Hložek et al., 2017).

In preclinical studies, dosing of THC is typically determined by whether it produces blood THC levels in the desired range seen in humans following cannabis consumption. However, whether route of administration influences how much THC, or its metabolites, accumulates in the brain and activates central cannabinoid CB1 receptors is not understood. As such, it is not clear if injections of THC that produce similar blood THC levels as those seen following inhalation are indeed comparable in the context of how much impact they have on activation of the central cannabinoid system. Consideration of the metabolism of THC is also incredibly important in this context and is often not measured in most analyses even though the metabolites of THC are highly bioactive, undoubtedly influenced by route of administration, and known to be significantly impacted by sex (Javadi-Paydar et al., 2017; Nguyen et al., 2020; Ruiz et al., 2021; Wiley et al., 2014). In the liver, THC is hydroxylated by cytochrome P450 enzymes into 11-hydroxy-THC (11-OH-THC), which is subsequently oxidized by the same group of enzymes to create 11-Nor-9-carboxy-THC (THC-COOH) and is excreted in urine (Huestis, 2005). The concentrations of THC metabolites are very important factors to consider when examining the impacts of THC administration, as 11-OH-THC is also psychoactive, is at least equipotent if not more potent than THC and diffuses more readily into the brain than THC (Grotenhermen, 2003; Lemberger et al., 1972; Perez-Reyes et al., 1972). Thus, differences in the generation and central accumulation of 11-OH-THC are not trivial and can have a robust impact on the outcome of studies given its ability to be as efficacious, if not more, as THC in activating CB1 receptors. THC-COOH is detectable for weeks, lacks any known psychoactivity, yet may possess anti-inflammatory and analgesic effects (Burstein, 1999; Grotenhermen, 2003).

In our attempts to develop more translationally relevant models of THC and cannabis administration, we examined if there were PK differences in the metabolism and accumulation of THC between inhaled and injected routes of administration that could help ascertain if these approaches are interchangeable or if there are differences that need to be considered when interpreting animal research data through a translational lens. To this extent, we utilized inhalation and injection paradigms, in both male and female rats, that produced comparable peak plasma THC concentrations to see if these different routes of administration resulted in differential PK or central accumulation of THC and its metabolites. To quantify concentrations of THC, 11-OH-THC and THC-COOH, we also developed our own analytical approach using mass spectrometry-liquid chromatography, which allowed us to quantify these molecules in both plasma and brain tissue.

## 2. Methods

### 2.1 Animals and housing

Adult male and female Sprague-Dawley rats were obtained from Charles Rivers Laboratories (St. Constant, QC, Canada). Rats were pair-housed with *ad libitum* access to water and standard laboratory chow and acclimated to a standard colony room (12 h light-dark cycle; constant temperature of 21 ± 1°C). Following ∼1 week of acclimation rats were split into two administration groups (injection [parenteral] and inhaled [pulmonary]) and each group was further sub-divided according to five timepoints (15, 30, 60, 90, and 240 min) (n=5-6 per group). All animal procedures were performed in accordance with the Canadian Council on Animal Care (CCAC) guidelines and were approved by the University of Calgary Animal Care Committee.

### 2.2 Injections of THC

Dosing for both injection and inhalation studies was based on pilot work establishing doses that produced roughly comparable blood THC levels in the range of 60-100 ng/mL. Rats received a single injection of THC intraperitoneally (dose of 2.5 mg/kg in a volume of 2 ml/kg). THC (100 mg/mL in 100% EtOH from Toronto Research Chemicals) was stored at -20°C until dissolved into a 1:1:8 solution of dimethylsulfoxide (DMSO), Tween 80, and 0.9% saline respectively. Following injections rats were euthanized via decapitation at five different timepoints (15, 30, 60, 90 and 240 min; referred to as INJ 15, INJ 30, INJ 60, INJ 90 and INJ 240 hereafter); trunk blood was collected and brains were extracted for hippocampus dissection. The hippocampus was chosen as the brain structure for analysis as it is an important site for many of the cognitive and emotional effects of cannabinoids, has a high density of cannabinoid receptors and is a brain structure whose isolation and dissection is consistent and straightforward. Blood samples were collected in EDTA tubes and stored on ice until centrifuged at 10,000 rpm for 10 minutes at 4°C. Plasma was collected and stored at -80°C until analysis. Dissected hippocampi were immediately frozen on dry ice and then stored at -80°C until analysis.

### 2.3 Passive inhaled delivery of THC

Rats received a single (15 min) session of inhaled exposure to a cannabis-derived THC-distillate (100 mg/mL) via a validated (Freels et al., 2020; Javadi-Paydar et al., 2017; Nguyen et al., 2016) vapour inhalation system (La Jolla Alcohol Research Inc., CA, USA). THC distillate (95% THC from Aphria Inc., ON, CND) was stored at room temperature until diluted to a concentration of 100 mg/mL THC in polyethylene glycol (PEG-400). The vapour inhalation system uses machinery similar to electronic cigarettes to deliver distinct “puffs” of cannabis vapour within airtight chambers. Chamber airflow is controlled by a vacuum compressor (i.e. a “pull” system) that draws room ambient air through an intake valve at a constant rate of 1 L per minute. At set intervals (as controlled by MedPC IV software [Med Associates, ST. Albans, VT, USA]) THC distillate is vaporized (utilizing a SMOK TFV8 X-baby atomizer [Shenzhen IVPS Technology Co., Shenzhen, China]) combining with the constant ambient air flow for delivery into the chamber. Air (and vapour) are evacuated through the back of the chamber via the vacuum compressor, filtered and ventilated out of the building (see Figure 1 for illustrated depiction of the vapour delivery system). In this study, THC vapor was delivered through a 10-second “puff” every 2 minutes for 15 minutes. In accordance with the injected animals, following the conclusion of the vapour session rats were euthanized via decapitation at five different timepoints (15 [immediate], 30, 60, 90 and 240 min, referred to as INH 15, INH 30, INH 60, INH 90 and INH 240 hereafter). Trunk blood and brains were collected and stored as previously described.

**Figure 1:**
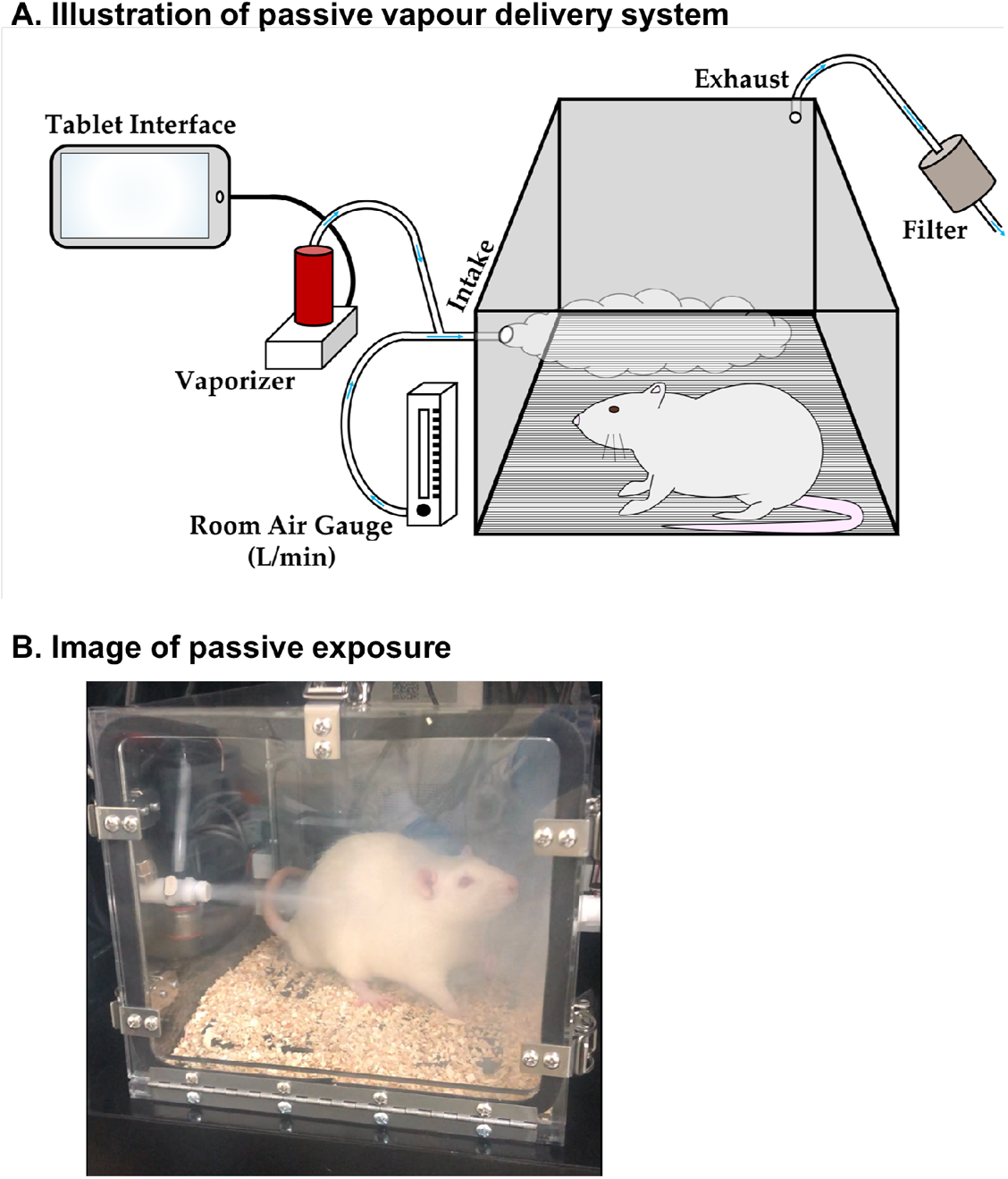
Vapor Delivery System. ***A. Illustration of passive vapour delivery system***: Schematic of vapour apparatus components with direction of air-flow (adapted from Freels et al. 2020). Briefly, the vapour inhalation system uses machinery similar to electronic cigarettes to deliver distinct “puffs” of vapour within airtight chambers. A vacuum compressor pulls ambient room air through an intake valve at a constant rate of 1 L per minute. At set intervals THC distillate is vaporized combining with the constant ambient air flow for delivery into the chamber. Air (and vapour) are evacuated through the back of the chamber via the vacuum compressor, filtered and ventilated out of the building. ***B. Image of passive exposure:*** Picture of a male SD rat within the vapour apparatus.

### 2.4 Body temperature

Body temperature was taken via a rectal thermometer immediately prior to euthanasia for all rats at the 30, 60, 90, and 240 min timepoints. As peak THC and metabolite levels were imperative to measure immediately following inhalation, blood and brain collection were prioritized at the 15 min timepoint. Further, hypothermic onset following THC exposure has been shown to occur at 30 min following inhalation (Manwell et al., 2014). Body temperature following THC administration was compared to control animals. Control animals were exposed to either vehicle vapour (PEG-400 alone) for 15 min or to vehicle injection (1:1:8 DMSO, Tween 80, and 0.9% saline) and their body temperature was taken at 30, 60, 90 and 240 min post administration.

### 2.5 Tandem mass spectrometry (LC-MS/MS)

#### 2.5.1 Standard solutions and reagents

Both standard and deuterated internal standard (IS) stock solutions were purchased from Cerilliant (Round Rock, TX, USA). The standard solutions, including Δ^9^-tetrahydrocannabinol (THC), 11-hydroxy-THC (11-OH-THC) and 11-nor-9-carboxy-THC (THC-COOH) were dissolved in acetonitrile at a concentration of 1.0 mg/mL. The IS stock solutions including THC-d3 and THC-COOH-d3 were dissolved in acetonitrile at 0.1 mg/mL. LC/MS grade acetonitrile, water and formic acid were purchased from Thermo Fisher Scientific (Edmonton, Canada). All compounds and their serial dilutions were stored at -80 °C freezer until use.

#### 2.5.2 Calibration curves

The stock solutions of each standard were mixed and diluted in 50% methanol/water to produce a set of standards ranging from 0.1 ng/mL to 100 ng/mL (0.1, 0.25, 0.5, 1, 2.5, 5, 10, 25, 50, 100). Internal standard (IS; d3 analytes) solution contains each compound at 10 ng/mL and was prepared in 50% methanol/water. Solutions used to establish calibration curves were prepared by mixing 20 µL of each standard solution and 20 µL of IS solution. The calibrators were analyzed in triplicate and the resulting calibration curves were fit by line of regression using a weight of 1/x^2^. R^2^ of each calibration curve was at least 0.999. Lower limit of quantitation (LLOQ) of each analyte was determined to be at 0.1 ng/mL.

#### 2.5.3 Sample preparation

Glass tubes containing 2 mL of acetonitrile and 100 µL of IS were prepared to receive plasma and brain samples. Each plasma sample was thawed at room temperature and 500 µL was directly pipetted into the prepared tubes. Each brain tissue sample was weighed and the frozen piece placed into the prepared glass tubes for manual homogenization with a glass rod until resembling sand. All samples were then sonicated in an ice bath for 30 min before being stored overnight at -20°C to precipitate proteins. The next day samples were centrifuged at 1800 rpm at 4°C for 3-4 min to remove particulates and the supernatant from each sample was transferred to a new glass tube. Tubes were then placed under nitrogen gas to evaporate. Following evaporation, the tube sidewalls were washed with 250 µL acetonitrile in order to recollect any adhering lipids and then again placed under nitrogen gas to evaporate. Following complete evaporation, the samples were re-suspended in 100 µL of 1:1 methanol and deionized water. Resuspended samples went through two rounds of centrifugation (15000 rpm at 4°C for 20 min) to remove particulates and the supernatant transferred to a glass vial with a glass insert. Samples were then stored at -80°C until analysis by LC-MS / Multiple Reaction Monitoring (MRM).

#### 2.5.4. LC-MS/MS analysis

LC-MS/MS analysis was performed using an Eksigent Micro LC200 coupled to an AB Sciex QTRAP 5500 mass spectrometry (AB Sciex, Ontario, Canada) at the Southern Alberta Mass Spectrometry (SAMS) facility located at the University of Calgary. The LC system consisted of a CTC refrigerated autosampler (held at 10 °C), a six-port sample injection valve with a 5 µL sample loop as well as a column oven. Chromatographic separation of the analytes was carried out on an Eksigent Halo C18 column (2.7 µm, 0.5 × 50 mm, 90Å, AB Sciex) at 40 °C. The mobile phase A was composed of 0.1% formic acid in water and the mobile phase B of 0.1% formic acid in acetonitrile. The analytes (2 µl injection) were eluted at 30 µl/min using a gradient from 25 to 95% B in 2.5 min. The column was then cleaned and regenerated using the following program: 95% B for 2 min, 95 to 25% B in 0.2 min and 25% B for 2.3 min. Before each injection, the column was equilibrated at initial LC condition for 1 min. Carryover was checked by injection of a blank in between samples. The data were acquired in positive electrospray ionization (ESI) and MRM mode. MRM transitions and collision energies (CE) of the different compounds are listed in Table 1. Each compound was acquired with two transitions. The first one was used to quantify the compound and the second one to discriminate isomers when necessary. Ion spray voltage was set at 5500 V. Nebulizer gas (GS 1), auxiliary gas (GS 2), curtain gas (CUR) were set at 30, 30, 35 (arbitrary units), respectively. Collision gas was set to Medium. Declustering potential (DP), entrance potential (EP) and cell exit potential (CXP) were set at 80, 7 and 14 V, respectively. LC-MS/MRM data were processed using Analyst 1.6 software (AB Sciex). Quantitation of each analyte was calculated using its extracted ion chromatogram (XIC; peak area) normalized by the peak area of its corresponding IS. Analyte concentration (in pmol/µL) were normalized to sample volume/weight and converted to ng/mL or ng/g for statistical analysis and graphing.

**Table 1:** Multiple Reaction Monitoring (MRM) Transitions and Collision Energies (CE) of different compounds/standards.

| **Compounds/Standards** | **Q1**<br>**(Da)** | **Q3**<br>**(Da)** | **Retention**<br>**Time (min)** | **CE**<br>**(volt)** |
| --- | --- | --- | --- | --- |
| 11-OH-THC-1 | 331 | 313 | 2 | 27 |
| 11-OH-THC-2 | 331 | 193 | 2 | 27 |
| THC-COOH-1 | 345 | 327 | 2 | 24 |
| THC-COOH-2 | 345 | 299 | 2 | 24 |
| THC-COOH-d3-1 | 348 | 330 | 2 | 24 |
| THC-COOH-d3-2 | 348 | 302 | 2 | 24 |
| THC-1 | 315 | 193 | 2.7 | 30 |
| THC-2 | 315 | 259 | 2.7 | 30 |
| THC-d3-1 | 318 | 196 | 2.7 | 30 |
| THC-d3-2 | 318 | 262 | 2.7 | 30 |

### 2.6 Statistical analysis

All data are expressed as mean ± SEM. Data were analyzed using IBM SPSS Statistics 26 and graphed using GraphPad Prism 8. Basal body temperature differed between males and females, so the two sexes were analyzed separately. Temperature data were analyzed by three-way ANOVA with drug group (THC and control), administration group (injection and inhalation), and timepoint (30, 60, 90, and 240 min) as between-subjects’ factors. Analyte data were analyzed by three-way ANOVA with sex (male and female), administration group (THC INH or INJ), and timepoint (15, 30, 60, 90 and 240 min) as between-subjects’ factors. Post-hoc comparisons used Bonferroni post hoc tests and differences were considered significant at p ≤ 0.05.

## 3. Results

### 3.1 Body Temperature

Body temperature measures were compared to controls and analyzed separately by sex (female > male, main effect of sex [F_(1, 136)_=23.219 at p<0.00001]). THC exposure resulted in hypothermia in male rats differentially depending on administration group (interaction effect of group and timepoint: F(_9,52_)=5.831 at p<0.0001, see Figure 2A). In particular, THC INH resulted in immediate hypothermia at 30 and 60 min (30 min: THC INH < CON INH, CON INJ, and THC INJ at p<0.05; 60 min: THC INH < CON INJ and THC INJ at p<0.05), whereas THC INJ resulted in delayed hypothermia at 90 and 240 min (90 min: THC INJ < CON INJ and THC INH at p<0.05; 240 min: THC INJ < CON INJ, CON INH, and THC INH at p<0.006). Along these lines, THC INH resulted in lower body temperature at 30 min compared to 90 and 240 min (p<0.005), as well as remained lower at 60 min compared to 90 min (p=0.014). In contrast, THC INJ resulted in lower body temperature at 90 min compared to 30 and 60 min (p<0.05), as well as remained lower at 240 min than all other timepoints (p<0.002).

**Figure 2:**
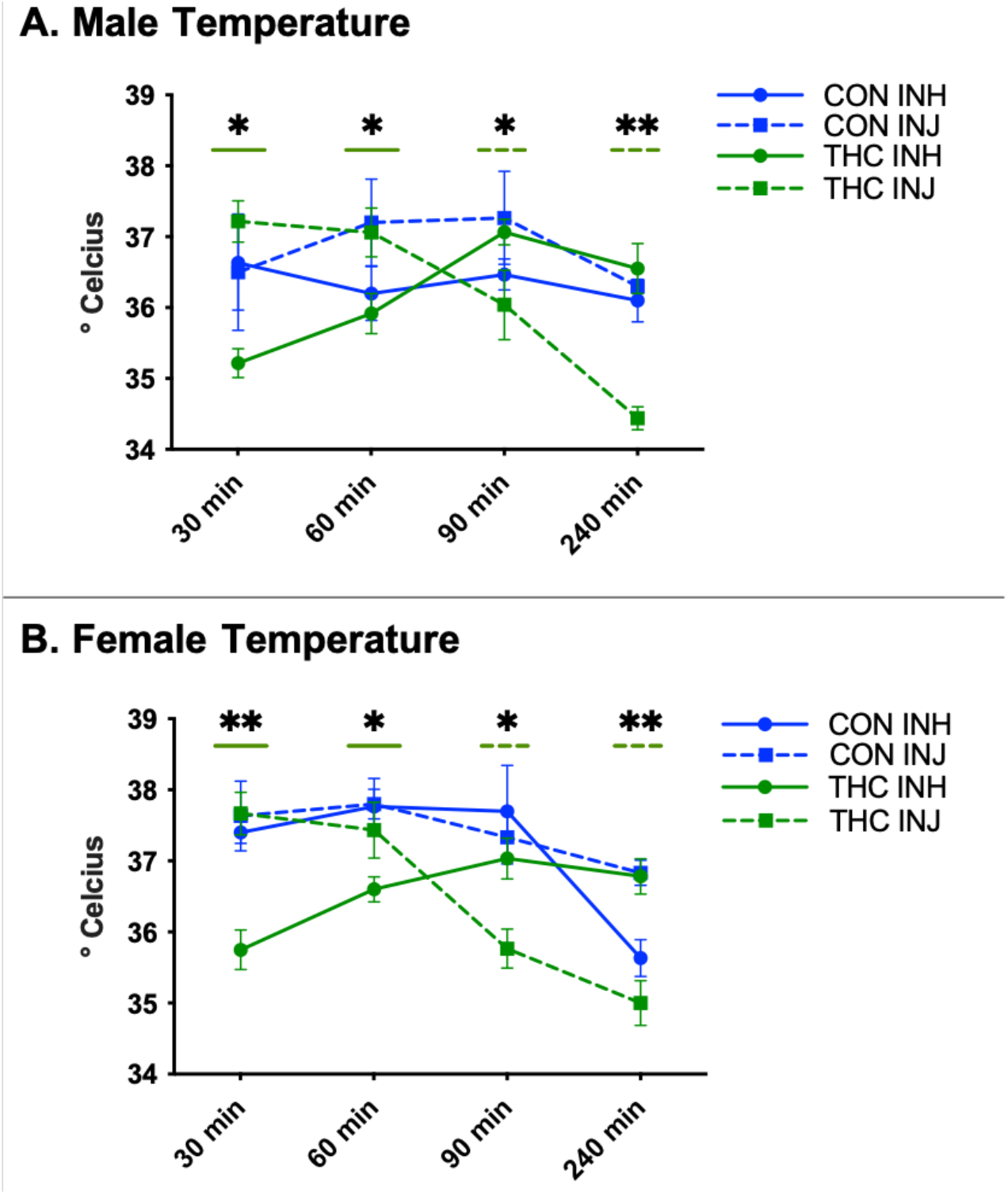
Body Temperature. ***A. Male temperature***: Data are presented as mean ± SEM; n=5-6 for each group. Presence of (*) with a solid and dashed green line indicates that THC INH and THC INJ respectively differ from one or more other groups. THC INH resulted in immediate hypothermia at 30 min (THC INH < all groups at *p<0.05) and at 60 min (THC INH < CON INJ and THC INJ at *p<0.05), whereas THC INJ resulted in delayed hypothermia at 90 min (THC INJ < CON INJ and THC INH at *p<0.05) and at 240 min (THC INJ < all groups at **p<0.01). ***B. Female temperature***: Data are presented as mean ± SEM; n=5-6 for each group. Presence of (*) with a solid and dashed green line indicates that THC INH and THC INJ respectively differ from one or more other groups. THC INH resulted in immediate hypothermia at 30 min (THC INH < all groups at **p<0.01) and at 60 min (THC INH < all groups at *p<0.05), whereas THC INJ resulted in delayed hypothermia at 90 min (THC INJ < all groups at *p<0.05) and at 240 min (THC INJ < CON INJ and THC INH at **p<0.01).

Females had an extremely similar pattern where THC exposure resulted in hypothermia in female rats differentially depending on administration group (interaction effect of group and time: F(_9,56_)=7.097 at p<0.00001, see Figure 2B). In particular, THC INH resulted in immediate hypothermia at 30 and 60 min (30 min: THC INH < CON INH, CON INJ, and THC INJ at p<0.001; 60 min: THC INH < CON INH, CON INJ and THC INJ at p<0.05), whereas THC INJ resulted in delayed hypothermia at 90 and 240 min (90 min: THC INJ < CON INJ, CON INH and THC INH at p<0.002; 240 min: THC INJ < CON INJ and THC INH at p<0.001). Along these lines, THC INH resulted in lower body temperature at 30 min compared to all other timepoints (p<0.05). Oppositely, THC INJ resulted in lower body temperature at 90 and 240 min compared to 30 and 60 min (p<0.001).

### 3.2 THC

Control values serve as assay controls and were undetectable. Analyte concentrations were therefore compared across THC administration groups and sex. THC chromatogram illustrates THC (black line) and THC-d3 (grey line) with an overlapping peak at 2.7 min (see Figure 3A).

**Figure 3:**
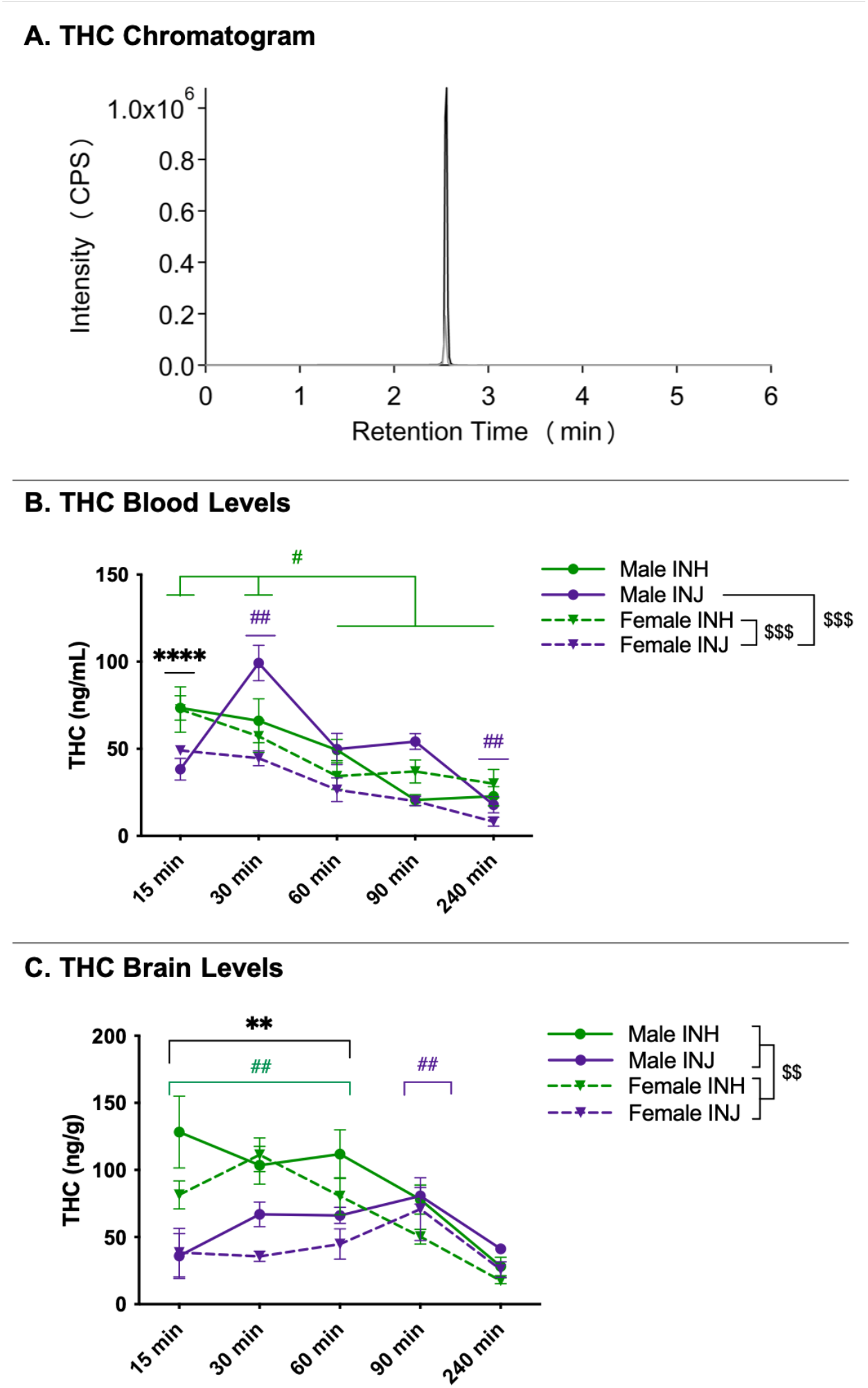
THC Chromatogram and Levels in Blood and Brain. ***A. LC-MS Chromatogram***: THC (black line) and THC-d3 (grey line) overlapping peaks at 2.7 min. ***B. Blood levels***: Data are presented as mean ± SEM; n=5-6 for each group. Presence of (*) indicates an administration difference with INH > INJ at 15 min at ****p<0.0001. Presence of (^#^) indicates a timepoint difference with green and purple lines indicating an INH and INJ difference respectively; specifically, INH 15 > all timepoints and INH 30 > INH 60/90/240 at ^#^p<0.05, whereas INJ 30 > all timepoints and INJ 240 < all timepoints at ^##^p<0.01. Presence of (^$^) indicates a sex difference; specifically, female INH > female INJ at ^$$$^p<0.001 and male INJ > female INJ at ^$$$^p<0.001). ***C. Brain levels:*** Data are presented as mean ± SEM; n=5-6 for each group. Presence of (*) indicates an administration difference with INH > INJ at 15, 30, and 60 min at **p<0.01. Presence of (^#^) indicates a timepoint difference with green and purple lines indicating an INH and INJ difference respectively; specifically, INH 15, 30, and 60 > INH 90 and 240 at ^##^p<0.01, whereas INJ 90 > INJ 15 and 240 at ^##^p<0.01. Presence of (^$^) indicates a sex difference with males > females at ^$$^p<0.01

In females, but not males, plasma THC concentrations were higher following INH than INJ regardless of timepoint (interaction effect of sex and group: F(_1,94_)=11.164 at p=0.0012, post hoc female INH > female INJ at p=0.0002, see Figure 3B). Following injection, but not inhalation, males had higher plasma THC concentrations than females regardless of timepoint (male INJ > female INJ at p=0.0002). Further, regardless of group, males and females differed in plasma THC concentrations across timepoints (interaction effect of sex and timepoint: F(_4,94_)=4.107 at p=0.004, post hoc male 30 > all timepoints at p>0.003, male 15 > male 90 at p=0.042, male 240 < male 15/30/60 at p<0.002, female 15 > all timepoints at p<0.02, female 30 > female 60/90/240 at p<0.02 see Figure 3B). Males also had higher plasma THC concentrations than females at 30 and 60 min (post hoc male 30 > female 30 at p=0.0005 and male 60 > female 60 at p=0.031). Finally, regardless of sex, plasma THC concentrations were higher following inhalation than injection at 15 min but not different at any other timepoint (interaction effect of group and timepoint: F(_4,94_)=5.301 at p=0.004, post hoc INH 15 > INJ 15 at p=0.00002, see Figure 3B). Plasma THC concentrations also differed by group across timepoint, such that following inhalation, plasma THC was higher at 15-min than all other timepoints and higher at 30-min than later timepoints (post hoc INH 15 > all timepoint at p<0.01 and INH 30 > INH 60/90/240 at p<0.03, see Figure 3B), whereas following injection, plasma THC was higher at 30-min compared to all other timepoints and lower at 240-min compared to all other timepoints (post hoc INJ 30 > all timepoints at p<0.002 and INJ 240 < all timepoints at p <0.008).

Brain THC concentrations were higher following INH than INJ at 15, 30 and 60 min regardless of sex (interaction effect of group and timepoint: F(_4,95_)=7.791 at p=0.00002; post-hoc INH 15 > INJ 15 at p<0.00001, INH 30 > INJ 30 at p=0.00004, and INH 60 > INJ 60 at p=0.003, see Figure 3C). Further, brain THC concentrations were higher at 15, 30 and 60 min compared to 90 and 240 min following inhalation (post-hoc INH 15 > INH 90 and 240 at p<0.003; INH 30 > INH 90 and 240 at p<0.001; INH 60 > INH 90 and 240 at p<0.01). Alternatively, brain THC concentrations were higher at 90 min than 15 and 240 min, and trending higher compared to 30 min, following injection (post-hoc INJ 90 > INJ 15 at p=0.006; INJ 90 > INJ 30 at p=0.07; INJ 90 > INJ 240 at p=0.003). Interestingly, brain THC concentrations were higher in males than females, regardless of group or timepoint (main effect of sex: F(_1,95_)=9.482 at p=0.003, see Figure 3C).

### 3.3 11-OH-THC

11-OH-THC chromatogram (see Figure 4A) illustrates 11-OH-THC (black line) and 11-OH-THC-d3 (grey line) with an overlapping peak at 2.0 min. Plasma 11-OH-THC concentrations were higher following INJ than INH at all timepoints regardless of sex (interaction effect of group and timepoint: F(_4,95_)=4.412 at p=0.003; post-hoc INH 15 < INJ 15 at p=0.03, INH 30 < INJ 30 at p<0.00001, INH 60 < INJ 60 at p<0.00001, INH 90 < INJ 90 at p<0.00001, INH 240 < INJ 240 at p=0.037). Further, while plasma 11-OH-THC concentrations did not differ across timepoint following inhalation, following injection plasma 11-OH-THC concentration were lower at 15 min compared to 30, 60, and 90 min (post-hoc INJ 15 < INJ 30 at p=0.0002, INJ 15 < INJ 60 at p=0.038, INJ 15 < INJ 90 at p=0.032), as well as lower at 240 min compared to 30, 60 and 90 min (post-hoc INJ 240 < INJ 30 at p<0.00001, INJ 240 < INJ 60 at p=0.0005, INJ 240 < INJ 90 at p=0.0005). Interestingly, plasma 11-OH-THC concentrations were higher in females than males, regardless of group or timepoint (main effect of sex: F(_1,95_)=4.613 at p=0.034, see Figure 4B).

**Figure 4:**
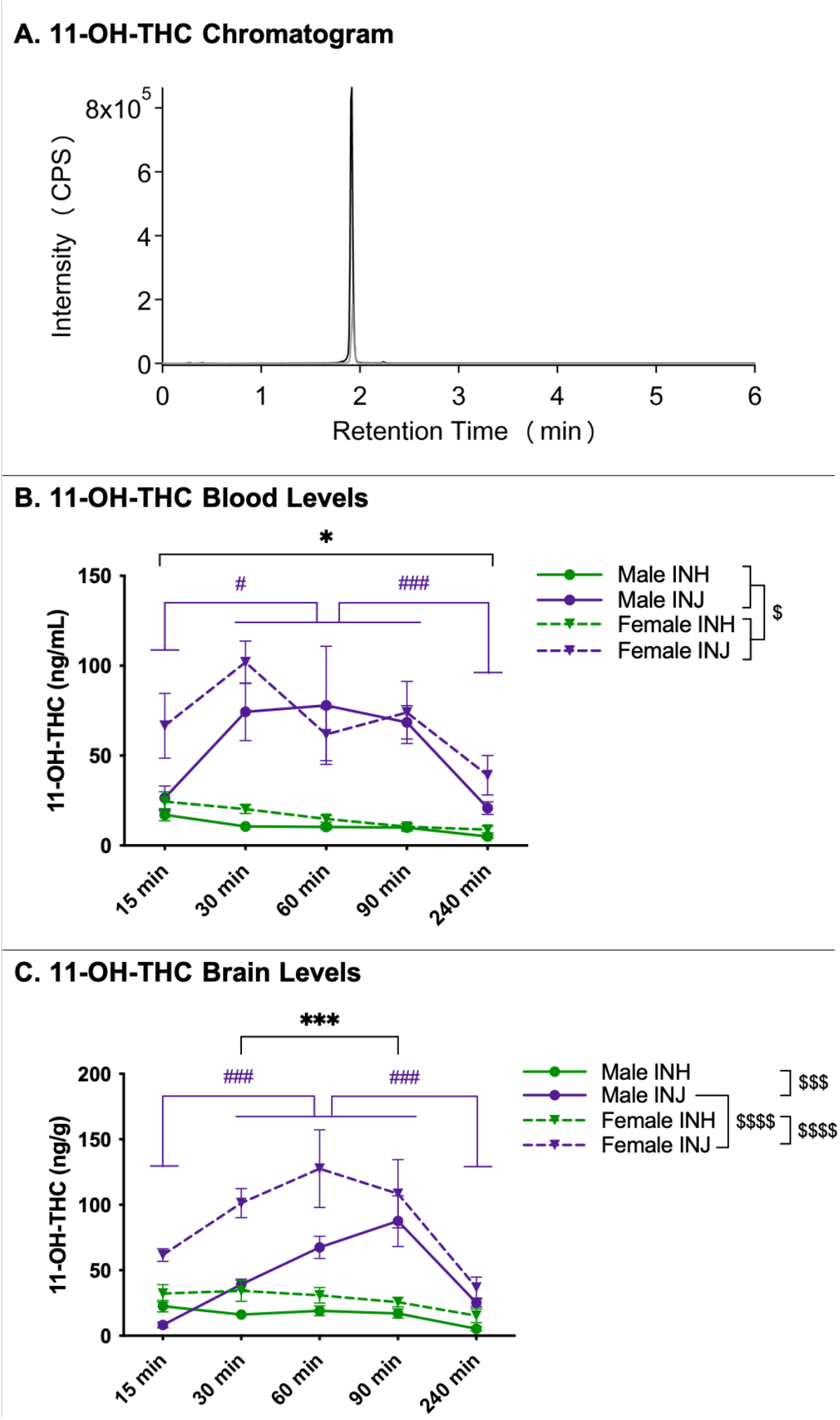
11-OH-THC Chromatogram and Levels in Blood and Brain. ***A. LC-MS Chromatogram***: 11-OH-THC (black line) and 11-OH-THC-d3 (grey line) peaks at 2.0 min. ***B. Blood levels:*** Data are presented as mean ± SEM; n=5-6 for each group. Presence of (*) indicates an administration difference with INH < INJ at all timepoints at *p<0.05. Presence of (^#^) indicates a timepoint difference with purple lines indicating an INJ difference where INJ 15 < INJ 30, 60, and 90 at ^#^p<0.05 and INJ 240 < INJ 30, 60, and 90 at ^###^p<0.001. Presence of (^$^) indicates a sex difference; specifically, females > males at ^$^p<0.05. ***C. Brain levels:*** Data are presented as mean ± SEM; n=5-6 for each group. Presence of (*) indicates an administration difference with INH < INJ at 30, 60, and 90 min at ***p<0.001. Presence of (^#^) indicates a timepoint difference with purple lines indicating an INJ difference where INJ 15 and 240 < INJ 30, 60, and 90 at ^###^p<0.001. Presence of (^$^) indicates a sex difference; specifically, male INH < male INJ at ^$$$^p<0.001, female INH < female INJ at ^$$$$^p<0.0001, and male INJ < female INJ at ^$$$$^p<0.0001.

In both sexes, brain 11-OH-THC concentrations were higher following INJ than INH regardless of timepoint (interaction effect of sex and group: F(_1,95_)=5.356 at p=0.023; post hoc male INH < INJ at p=0.0002 and female INH < INJ at p<0.00001, see Figure 4C). Further, following injection, brain 11-OH-THC concentration were higher in females than males (post hoc male INJ < female INJ at p<0.00001). Interestingly, regardless of sex, brain 11-OH-THC concentrations were higher following INJ than INH at 30, 60 and 90 min (interaction effect of group and timepoint: F(_4,95_)=6.411 at p=0.0001; post hoc INH 30 < INJ 30 at p=0.0002, INH 60 < INJ 60 at p<0.00001, INH 90 < INJ 90 at p<0.00001, see Figure 4C). Further, while brain 11-OH-THC concentrations did not differ across timepoint following inhalation, following injection brain 11-OH-THC concentration were lower at 15 min compared to 30, 60, and 90 min (post-hoc INJ 15 < INJ 30 at p=0.0009, INJ 15 < INJ 60 at p<0.00001, INJ 15 < INJ 90 at p<0.00001), as well as lower at 30 min compared to 60 min (post hoc INJ 30 < INJ 60 at p=0.023) and at 240 min compared to 30, 60 and 90 min (post-hoc INJ 240 < INJ 30 at p=0.001, INJ 240 < INJ 60 at p<0.00001, INJ 240 < INJ 90 at p<0.00001).

### 3.4 THC-COOH

THC-COOH chromatogram (see Figure 5A) illustrates THC-COOH (black line) and THC-COOH-d3 (grey line) with an overlapping peak at 2.0 min. In both sexes, plasma THC-COOH concentrations were higher following INJ than INH regardless of timepoint (interaction effect of sex and group: F(_1,93_)=12.241 at p=0.0007; post hoc male INH < INJ at p=0.00006 and female INH < INJ at p<0.00001, see Figure 5B). Further, following injection, plasma THC-COOH concentration were higher in females than males (post hoc male INJ < female INJ at p<0.00001). Interestingly, regardless of sex, plasma THC-COOH concentrations were higher following INJ than INH at 30, 60, 90 and 240 min (interaction effect of group and timepoint: F(_4,93_)=3.0 at p=0.022; post hoc INH 30 < INJ 30 at p<0.00001, INH 60 < INJ 60 at p<0.00001, INH 90 < INJ 90 at p<0.00001, INH 240 < INJ 240 at p=0.00003, see Figure 5B). Further, while plasma THC-COOH concentrations did not differ across timepoint following inhalation, following injection plasma THC-COOH concentration were lower at 15 min compared to all other timepoints (post-hoc INJ 15 < all timepoints at p<0.003).

**Figure 5:**
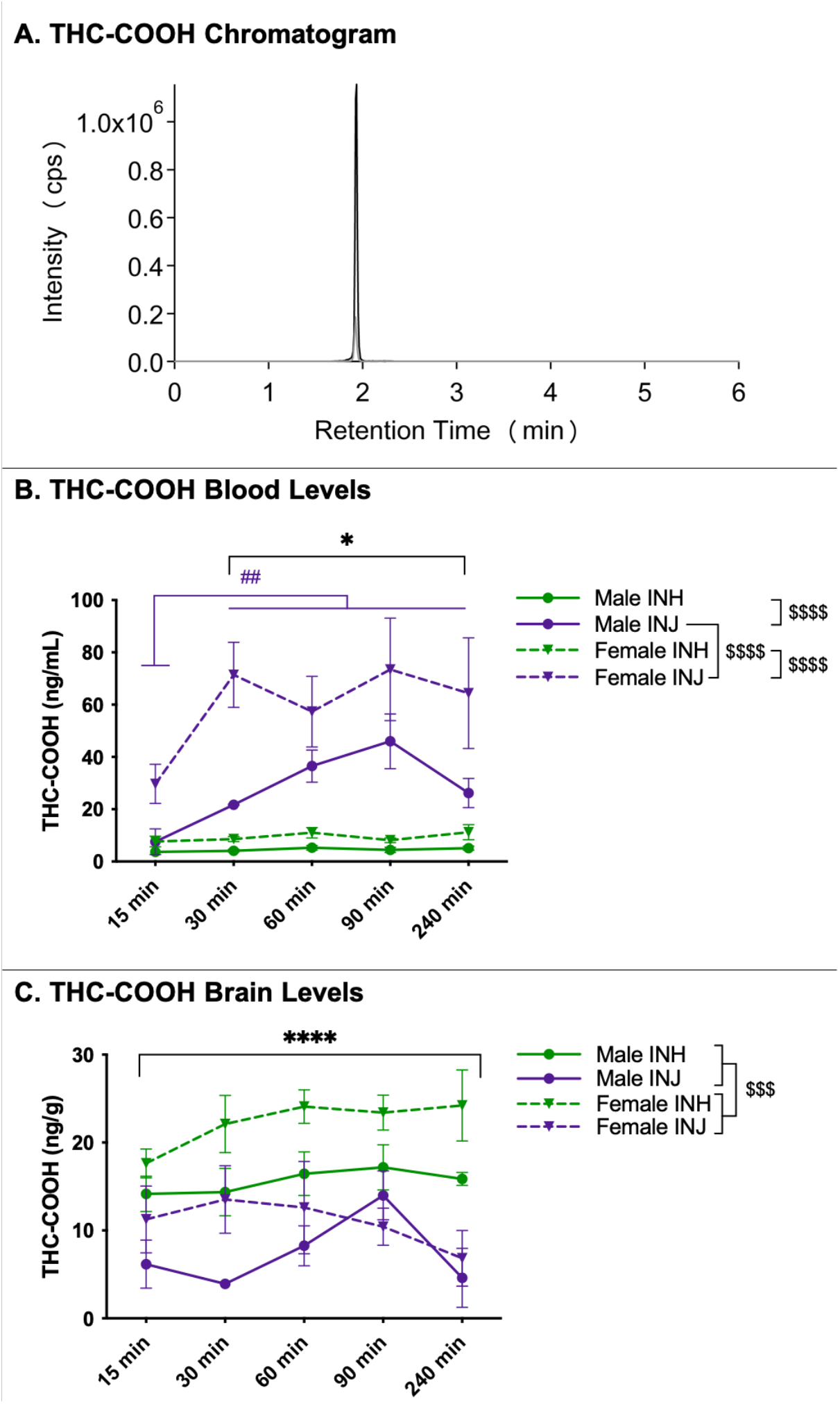
THC-COOH Chromatogram and Levels in Blood and Brain. ***A. LC-MS Chromatogram***: THC-COOH (black line) and THC-COOH-d3 (grey line) peaks at 2.0 min. ***B. Blood levels:*** Data are presented as mean ± SEM; n=5-6 for each group. Presence of (*) indicates an administration difference with INH < INJ at 30, 60, 90 and 240 min at *p<0.05. Presence of (^#^) indicates a timepoint difference with purple lines indicating an INJ difference where INJ 15 < all timepoints at ^##^p<0.01. Presence of (^$^) indicates a sex difference; specifically, male INH < male INJ and female INH < female INJ at ^$$$$^p<0.0001, and male INJ < female INJ at ^$$$$^p<0.0001. ***C. Brain levels:*** Data are presented as mean ± SEM; n=5-6 for each group. Presence of (*) indicates an administration difference with INH > INJ at ****p<0.0001. Presence of (^$^) indicates a sex difference with males < females at ^$$$^p<0.001.

Brain THC-COOH concentrations were higher in females than male regardless of group or timepoint (main effect of sex: F(_1,95_)=15.442 at p=0.0002, see Figure 5C). Further, brain THC-COOH concentrations were higher following inhalation than injection regardless of sex or timepoint (main effect of group: F(_1,95_)=56.656 at p<0.00001, see Figure 5C).

## 4. Discussion

Given the increased usage, demand, and potency of cannabis over the last few years (ElSohly et al., 2016; Government of Canada, 2019; Goodman et al., 2020), establishing an animal model that closely reflects human consumption is critical in order to study the impact on brain and behaviour.

Nevertheless, most animal studies to date examine the effects of cannabis through IP injection of THC, which produces markedly different behavioural effects than inhalation (Manwell et al., 2014). In efforts to make THC injection studies possess more face validity for translatability to humans, these previous studies aimed to produce peak plasma THC concentrations that are comparable to concentrations in human cannabis smokers. Utilizing ‘comparable’ dosages (based on their ability to produce blood THC levels in the range produced in humans from cannabis) established in previous literature (Javadi-Paydar et al., 2017; Nguyen et al., 2016) as well as pilot work in our laboratory, we sought to produce similar peak plasma THC concentrations following inhalation and injection that are within the range typically seen following cannabis consumption in humans (Abrams et al., 2007; Hartman et al., 2015; Huestis et al., 1992; Huestis and Cone, 2004; Newmeyer et al., 2016; Schwope et al., 2011), to determine if route of administration influenced metabolism or central accumulation of THC. Our data provide clear evidence on the different physiological response and PK profiles of THC and its metabolites following comparable dosing of inhaled versus injected THC, which could have significant impacts for data interpretation and generalizability. Importantly, we found that inhalation led to immediate hypothermia and an initial higher plasma and brain THC concentration, while injection led to delayed hypothermia, dramatically higher 11-OH-THC concentrations in both plasma and brain and higher THC-COOH concentration in the brain. Males in general had higher THC levels while females had higher metabolite levels, supporting previous work that there are robust sex differences in the PK of THC.

### Body Temperature

It is well established that THC exposure induces a hypothermic response in both humans and animals (Huestis, 2007; Javadi-Paydar et al., 2017; Nguyen et al., 2020).

Hypothermia was found in both males and females following inhalation and injection of THC. In accordance with previous literature (Javadi-Paydar et al., 2017; Nguyen et al., 2020), inhalation led to immediate hypothermia with lower temperatures at 30 and 60 min following THC administration which normalized by 90 and 240 min, whereas injection results in the opposite effect of delayed hypothermia with lower temperatures at 90 and 240 min compared to 30 and 60 min following THC administration. This hypothermic difference based on route of administration is not surprising given that inhaled THC quickly enters the bloodstream and is taken up by the brain, whereas injected THC undergoes metabolism by the liver before reaching the systemic circulation (Turner et al., 2011). Along these lines, the onset of the hypothermic response seen in this study occurs in conjunction with peak brain THC levels following both inhalation and injection, at 30 and 90 min respectively, indicating that this is a good physiological proxy readout of central accumulation of cannabinoids. The impact of route of administration on the temporal kinetics of hypothermia was not influenced by sex.

### THC Concentrations in Plasma and Brain

Utilizing ‘comparable’ doses of THC inhalation and injection, as determined by previous literature and our pilot work (Javadi-Paydar et al., 2017; Nguyen et al., 2016), we show altered plasma THC levels across route of administration and timepoint.

Specifically, inhalation led to higher plasma THC concentrations than injection at 15 min, but the two routes of administration did not differ at any other timepoint. This is not surprising as inhalation results in much more rapid uptake into the bloodstream than injection. In fact, it is known that inhalation produces peak plasma THC levels 10-15 min after initial administration in humans and rodents (Hartman et al., 2015; Hložek et al., 2017; Huestis and Cone, 2004; Newmeyer et al., 2016; Schwope et al., 2011), whereas peak levels are found at a slightly later timepoint following injection in rodents (Javadi-Paydar et al., 2017; Nguyen et al., 2016; Taffe et al., 2020; Torrens et al., 2020). Along these lines, plasma THC concentrations were highest following inhalation at 15 and 30 min compared to 60, 90 and 240 min. Alternatively, plasma THC concentrations were higher following injection at 30 min compared to all other timepoints. As anticipated, there was no difference between peak THC levels between the two routes of administration (peak THC following inhalation 73 ng/mL vs. peak THC following injection 72 ng/mL), and importantly these levels fall within the range found in human studies (60-200 ng/mL) (Abrams et al., 2007; Hartman et al., 2015; Huestis et al., 1992; Huestis and Cone, 2004; Newmeyer et al., 2016; Schwope et al., 2011).

Despite the similar peak plasma THC concentrations, inhalation led to higher brain THC concentrations at 15, 30, and 60 min compared to injection. This is not surprising as cannabis inhalation provides rapid delivery into the blood stream, bypasses initial liver metabolism, and results in more immediate uptake by highly perfused tissues, such as the brain (Huestis, 2007, 2005; Turner et al., 2011). Further, in accordance with previous literature showing earlier peak brain THC concentrations following inhalation than injection (Hložek et al., 2017), we found brain THC levels peaked at 15 and 30 min following inhalation, while peak brain THC levels did not occur until 90 min following injection. Interestingly, the peak subjective “high” in humans following cannabis inhalation is ∼30 min following onset of administration (Grotenhermen, 2003), which corresponds to the higher THC concentrations in the brain in this study following inhalation.

### 11-OH-THC Levels in Plasma and Brain

Overall, injection administration yielded significantly higher plasma and brain concentrations of 11-OH-THC compared to inhalation. Specifically, both plasma and brain concentrations of 11-OH-THC were relatively low (∼13 and ∼22 ng/mL respectively) following inhalation and did not differ across timepoints. Low concentrations of 11-OH-THC following inhalation are common as concentrations are recirculated through the enterohepatic pathway (i.e. liver to bile to small intestine back to liver) and quickly metabolized to THC-COOH (Grotenhermen, 2003). Alternatively, plasma concentrations of 11-OH-THC were highest at 30, 60, and 90 min compared to 15 and 240 min following injection, reaching average peak levels of ∼88 ng/mL, which is about 8 times higher than inhalation levels. Further, brain concentrations of 11-OH-THC were also highest at 30, 60, and 90 min compared to 15 and 240 min following injection, reaching average peak levels of ∼98 ng/g, which is about 4.5 times higher than inhalation levels. This striking difference in the production of 11-OH-THC is not trivial because 11-OH-THC is an agonist at CB1 receptors, is psychoactive, is believed to pass into the brain more readily than THC (which is consistent with the difference in blood-brain ratios seen for THC vs 11-OH-THC), and is as, or more, potent than THC in its ability to produce behavioural and physiological effects (Grotenhermen, 2003; Lemberger et al., 1972; Perez-Reyes et al., 1972). Recognizing that it is ultimately the activation of CB1 receptors in the brain, which is readily achieved by both THC and possibly more so by 11-OH-THC, the overall impact administration of THC will have on central CB1 receptor activation must take both THC and 11-OH-THC levels into account. Following injection, extremely high concentrations of 11-OH-THC in the brain will produce different physiological, psychological, and behavioural effects as compared to inhalation. In fact, given the dramatic accumulation of 11-OH-THC in the brain following injection, as well as its significant potency at the CB1 receptor, it seems reasonable to hypothesize that injection of THC produces a much more robust and sustained activation of brain CB1 receptors than THC administered via inhalation. As our data indicate that inhalation produces a rapid accumulation of THC in the brain, followed by relatively rapid clearance, this would suggest that the ability of inhaled THC to activate brain CB1 receptors is likely a time-limited effect. This is consistent with our hypothermia data and the relatively rapid peak, and diminution, of psychological effects and intoxication produced by inhaled cannabis or THC. Alternatively, injected THC produces lower initial brain THC concentrations compared to inhaled with levels accumulating over time to reach peak THC levels at 90 min that are comparable to inhaled. However, injection administration also includes the addition of high 11-OH-THC levels, produced through hepatic metabolism, and sequestered into the brain. As such, injections of THC will potentially produce profoundly different biological effects since the magnitude of CB1 receptor activation (through brain levels of both THC and 11-OH-THC) will be much greater than that following inhaled THC. Given that there is a notable discrepancy between many of the beneficial and adverse impacts of THC that have been documented in rodent studies using injection-based approaches relative to human studies examining cannabis users, one must consider that the injection-based approach for THC has limitations for translational research. As the cellular impacts of CB1 receptor activation will be influenced by the magnitude and duration of its activation, the impacts of accumulating 11-OH-THC in the brain following injections of THC must be considered in future rodent studies.

### THC-COOH Levels in Plasma and Brain

Injection administration yielded higher levels of plasma THC-COOH but lower levels of brain THC-COOH compared to inhalation. More specifically, plasma concentrations of THC-COOH were relatively low (∼7 ng/mL) following inhalation and did not differ across timepoints. Whereas, plasma concentrations of THC-COOH were highest at 30, 60, 90 and 240 min compared to 15 min, reaching average peak levels of ∼60 ng/mL, about 9 times higher than inhalation levels. Along these lines, previous studies have also shown peak THC-COOH concentrations at later timepoints (60-120 min) following injection in rats (Torrens et al., 2020) and inhalation in humans (Huestis et al., 1992). Further, given that plasma 11-OH-THC concentrations were higher following injection, and 11-OH-THC is the metabolic precursor of THC-COOH, it is not surprising that THC-COOH follows the same pattern. Low levels of THC-COOH in the brain are anticipated as it is the primary metabolite for urinary elimination (Huestis, 2005).

### Sex Differences

Females exhibited significantly higher levels of 11-OH-THC and THC-COOH in both brain and blood, indicating that females metabolize THC at a faster rate than males do, which is relatively consistent with previous work (Ruiz et al., 2021; Wiley et al., 2014). In line with accelerated metabolism of THC and increased concentrations of THC metabolites in females, females were found to have lower levels of THC itself relative to males, particularly in the brain. Previous studies have shown no differences between male and female THC blood and brain levels (Javadi-Paydar et al., 2017; Nguyen et al., 2020; Ruiz et al., 2021; Wiley et al., 2014). This sex difference in the production of 11-OH-THC, particularly following injections, has relevance for interpreting preclinical studies. For example, many studies report that females are more sensitive to the effects of THC relative to males, particularly in the context of adverse effects of THC. Our data suggests that while females achieve slightly lower brain THC levels than males, they appear to acquire brain 11-OH-THC levels that are essentially double that seen in males following injection of THC. Given the bioactivity and potency of 11-OH-THC, this suggests that injection-based studies of THC may suggest sex differences exist in some outcomes, but this effect may be an artifact of the elevated levels of 11-OH-THC produced by injection; an effect which does not occur in the same manner following inhalation where brain 11-OH-THC levels are quite low and comparable between males and females. These sex differences in THC metabolism may also have implications for human THC consumption, especially when it is consumed via an oral route as entero-hepatic metabolism will impact THC metabolism.

### Limitations and Conclusions

Of note, the current studies do not include oral or edible intake of cannabis, another popular form of consumption. This is an area of future work in our group and has recently been successfully executed by others (Kruse et al., 2019). Oral consumption would also result in an increase in 11-OH-THC production due to first pass metabolism; however, given that an intoxicating dose of oral cannabis in humans produces peak blood levels of THC in the range of 1-5 ng/m (Vandrey et al., 2017), which is about 1/20-1/100 of the current level produced by injection of THC, one can anticipate that the levels of 11-OH-THC produced would be substantially lower then what we see following injection. Thus, despite the potential similarities in pharmacokinetic trajectories, injections would not be a suitable comparison for oral routes of administration of cannabis or THC.

In conclusion, our data demonstrate significant and biologically relevant differences in the pharmacokinetics and accumulation of THC and its metabolites following injection versus inhalation. The current study generally supports previous findings but provides the first direct comparison of both sex and route of administration of THC to reveal an accurate picture of how these variables are impacting THC metabolism and central accumulation. These findings should be considered for translational preclinical studies attempting to model the impacts of cannabis or THC on the brain and behavioural processes. IP injections are the most frequent route of administration for animal models examining the effects of cannabis (THC) and many previous studies claim the translatability and relevance to human consumption by aiming to produce peak plasma THC concentrations that are comparable to concentrations in human cannabis smokers. Utilizing doses that produced comparable peak plasma THC concentrations, our study illustrates robust differences in the pharmacokinetics and central accumulation of THC and its bioactive metabolites when administered via injection versus inhalation. These differences likely underlie the inconsistency of reproducible behavioural findings between THC injections and inhalation, and support the importance of appropriately modeling the route of administration in preclinical studies. This is not uncommon in the drug research field, and in fact studies utilizing injections of ethanol have long been abandoned over the appropriate use of ethanol vapour or drinking as this produces comparable effects to humans, and also allows for the study of volitional administration. Based on the data generated herein, we suggest that researchers conducting translational work in the realm of THC and cannabis strongly consider utilizing inhalation models, or oral routes of administration, to ensure that any biological effects they see from THC or cannabis extract administration are not artifacts produced by the accumulation of 11-OH-THC in the brain and activating CB1 receptors in a temporal manner that is likely quite distinct from what is occurring with humans during typical cannabis use.

## Acknowledgements

Thank you to all current and former members of the Hill laboratory for their assistance and expertise. Thank you to Maury Cole and La Jolla Alcohol Research Inc. for his technical assistance with the vapor chambers and Cece Hillard for her input on the manuscript.

## Funding

This study was supported by operating funds from the Canadian Institutes of Health Research (CIHR TCP-431510; MNH) and the Hotchkiss Brain Institute (MNH and SB). CH received salary support from Eyes High Postdoctoral Scholarship and Harley Hotchkiss – Samuel Weiss Postdoctoral Fellowship. SLB received salary support from a Vanier Scholarship from CIHR, GNP received salary support from the Branch Out Neurological Foundation (BONF), RJA received salary support from the Mathison Centre for Mental Health Research & Education, SHML received salary support from the William H. Davies Medical Research Scholarship. All funding agencies had no influence on the design, execution, or publishing of this work.

## Conflict of Interest Statement

MNH is a member of the scientific advisory board for Shoppers Drug Mart, Jazz Pharmaceuticals and Lundbeck.

